# Identification of Banana heat responsive long non-coding RNAs and their gene expression analysis

**DOI:** 10.1101/2022.05.09.491188

**Authors:** Vidya Srinivasa Murthy, Swarupa Venkatesa Murthy, Pooja Ramesh, Vidya Niranjan, Vasanth Kumar, Vidya Bharthi H, Ravindra M. Bhatt, Laxman R. Hunashikatti, Kundapura Venkataramana Ravishankar

**Author notes:** Corresponding author: Vidya Srinivasa Murthy, E-mail addresses.

## Abstract

Identification and characterization of long non-coding RNAs (lncRNAs) in the last decade has attained great attention because of their importance in gene regulatory functions in response to various plant stresses. Rising temperature is a potential threat to the agriculture world. Banana being an important economic crop, identification and characterization of genes and RNAs that regulate high temperature stress is imperative. As the prediction of lncRNAs in response to heat stress in banana is largely unknown, the present study was focused to identify the heat stress responsive lncRNA in banana (DH Pahang). Perl script was written to identify the novel transcripts, using the work flow. StringTie and CPC softwares were used and a total of 363 novel HS-lncRNA were identified in banana. Further, lncRNA were classified as 288 lincRNAs, 71 antisense LncRNA, 5 sense lncRNAs. The functional classification was done and transcripts were broadly classified into molecular function, cellular components and biological processes. Differential expression of lncRNA showed the varied patterns at different stages of heat stress. Finally, qPCR results confirmed DGE expression pattern of lncRNAs. Further, the Cytoscape analysis was performed which showed protein coding genes involved in membrane integrity and other signal transduction pathways. Taken together, these findings expand our understanding of lncRNAs as ubiquitous regulators under heat stress conditions in banana.

## Introduction

Abiotic stress, like extreme heat, salinity, drought affects crop growth and productivity. Plants have evolved through these environmental changes occurring across the globe over the years. Interestingly, heat stress due to climate change has gained importance in recent times. Many studies on identification of stress responsive genes and regulatory pathways involve heat shock proteins (HSP) and heat shock factors (HSF) during thermotolerance (Vidya et al. 2017). Banana being a tropical crop, it experiences high temperature stress during its growth period, affecting productivity and products. The most effective way to overcome this problem is to improve the thermotolerance of the banana cultivars. However, breeding programme remains a little complicated for banana due to its genetic makeup and sterility. Hence, it is necessary to understand regulatory mechanism operating in banana especially gene regulation including involvement of lncRNA.

As a result, with the development of transcriptome sequencing and computational methods, systematic identification and classification of lncRNAs has been carried out in a variety of species. lncRNA have a basic classification where they are further classified into microRNA (miRNA), circular RNA (circRNA), long non coding RNA (lncRNA), and small interfering RNA (siRNA). LncRNAs are arbitrarily defined as a set of non-coding RNA those consist of a length greater than 200 nucleotide length. They are further classified based on the loci positions like long intergenic non coding RNA (lincRNA), antisense lncRNA, intronic lncRNA, overlapping lncRNA (Zhang et al. 2014). They are known to regulate and involve in abiotic and biotic stress tolerance mechanism. However, information on lncRNA under heat stress is completely lacking in banana. Keeping this in view, the present study envisages to identify lncRNA in banana during heat stress and further validation using qPCR. The results will help us in understanding role of the HS-LncRNA in thermotolerance.

## Material and Methods

### Plant material

Banana tissue cultured plants (5-6 weeks old). cv Grand Naine was used in this study. As a part of temperature induction response (TIR) analysis were imposed with mild heat stress referred as Induction stress (I) : Gradual increase in temperature from 30ºC to 42 ºC for 2.5 hours, Induction+lethal (I+L): Plants after induction (from 30ºC to 42 ºC for 2.5 hours), were shifted to 55 ºC for 2hours, Lethal: Un-induced, control plants, directly shifted to 55 ºC and kept for 2hours (Vidya et al. 2017).

### Data sets

RNA was isolated and transcriptome analysis was performed using illumina platform in M/S Genotypic Pvt Ltd, Bengaluru facility. We used four libraries 1) control 2) Induction (I) 3) Induction followed by lethal (I+L) 4) Lethal (L). Banana genomic sequences, CDSs were downloaded from the link (http://banana-genome.cirad.fr). Our transcriptome data were deposited at NCBI with SRA number (**SRP074337:PRJNA320030**). These four libraries were used to mine lncRNA and pipeline was developed to identify putative lncRNAs (Figure 1) (Vidya et al. 2018)

**Figure 1:**
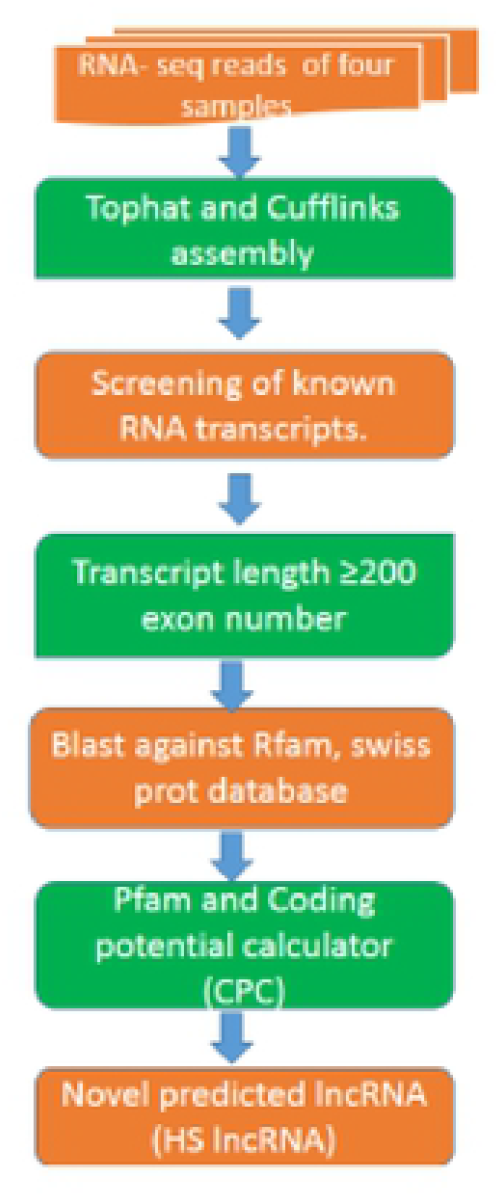
The flowchart for long non-coding RNA prediction responsive for heat stress.

### Bioinformatic analysis

The reads obtained from each library were mapped to the banana reference genome (*M. acuminata* ssp. *malaccensis* var. Pahang (DH Pahang) using Tophat 2.0 (Trapnell et al. 2009). The uniquely aligned reads using Cufflinks (version 2.0.1) transcriptome were assembled separately (Trapnell et al. 2010). A non-redundant genome annotation file was obtained by aligning with the banana reference genome, which is called reference annotation based transcript assembly.

Database building, of the annotated transcripts UniRef (UniProt Consortium, 2013) was used. In UniRef, the database is built using TrEMBL and SwissProt headers where the protein id, taxonomy, names are generated. Cuff merge programme was run where the transcripts were merged into a unified list. Using this merged file as an input (.gtf) the cuff diff compares between each sample and also its isoforms.

### Identification of LncRNA

Similarity search for lncRNA was done using BLASTX software. This results was then compared against UniRef and SwissProt. BLASTN was conducted to improve the confidence of the lncRNA which was run against Rfam and rRNA. Here the protein coding genes are excluded, and in further step where dna2pep (Wernersson et al. 2006) and Portrait software (Arrial et al. 2009) were used to identify the longest ORFs (<100aa & >200nt) and non-coding potential transcripts. Coding potential calculator software (CPC) was used (http://cpc.cbi.pku.edu.cn) (Kong et al. 2007) after stringent filtering criteria with < -1 CPC values were considered for further analysis. The transcripts were further aligned to NCBI protein database and those transcripts with known protein domains were excluded based on Pfam HMM (http://pfam.sanger.ac.uk/) (Bateman et al. 2004) using HMMER 3.0 software (HMMER 3.0) (http://hmmer.janelia.org/) with default parameters. Finally, 363 lncRNA were identified which were later classified into categories like long intergenic non coding RNAs (lincRNA), intronic lncRNA, anti-sense lncRNA.

### Circos and Cytoscape tool

Circos is a tool used for identification of the location of the genes or lncRNAs and similarities and differences arising on a given chromosome and its distribution in the genome (Krzywinski et al. 2009). This tool uses a layout like circular ideogram which helps in drawing a related pattern between the position and its pair with the help of ribbons. Since it’s a user friendly tool, it allows to flexibility and rearrangements of different elements in the image obtained. The venn diagrams were built using an online available tool (http://bioinfogp.cnb.csic.es/tools/venny/).

Cytoscape is an open source software which aids in developing a complex biological network, bio-molecular interaction with different states such as phenotypic data or expression profiles and therefore link the data through functional annotation. In our study, the plug-in used to build a co-expression pattern was with 100kb space (http://www.cytoscape.org/) (Cline et al. 2007).

### Validation using quantitative real-time PCR analysis

To elucidate the validity of the data qPCR analysis was performed. cDNA was synthesised from 1 μg of total RNA using reverse transcription Kit (#K1622, Thermo Scientific) following manufacturer’s instructions. Further, this template was employed for qPCR using DyNAmo Flash SYBR Green qPCR kit (#F-416L, Thermo Scientific) and lncRNA specific primers. *GAPDH* was used as an internal control (Swarupa et al. 2014). Primers used in this study are listed in table S1. Each reaction (20 μL) consisted of a 2ng of cDNA, 1μL of each primer (5pico moles concentration) and 10μL of SYBR mix. Thermal cycling conditions were 40 cycles of 95 °C for 15s, 60 °C for 30s (data collection point) and 72 °C for 15s followed by dissociation curve. All the reactions were performed for three biological replicates and with three technical replicates. The comparative CT method (2^-ΔΔct^) was used to quantify the relative expression of lncRNA (Bustin et al. 2004).

### Statistical analysis

The data was analyzed statistically using Microsoft Excel (2010). All values were presented as mean ± standard deviation. A value P<0.05 was considered to be statistically significant.

## Results and Discussion

### Genome wide identification of lncRNAs

RNA-seq, paired end sequencing was performed (SRP074337:PRJNA320030) and the sequences were mapped to the *Musa acuminata* (DH Pahang version 2) (Martin et al. 2016). The details and the work flow, followed are described in the Figure 1. TopHat2 was used to align the transcripts to the reference genome. StringTie was used in the next step which produced more accurate and complete transcriptome assembly and was found to run faster over the data sets compared to other software like cuffinks (Pertea et al. 2015). Step 3, FEELnc a module was used which aims in identifying LncRNA from the .gtf file obtained in the second step. 3a) FEELnc filter, this module helps us to filter out the short reads i. e. >200 nucleotides. 3b) FEELnc coding potential module (FEELnc codpot), this module computes the CPS and annotates the ORFs. 3c) FEELnc classifier module (FEELnc classifier), the classification is based on the type, direction, location. Additionally, it also can be used to classify transcripts based on biotypes (snoRNAs). Step 4, the number of LncRNA obtained from the above process was taken further to Annocript, where different data base information about the known or annotated transcript were obtained. After this, in the Step 5, the un-annotated transcripts were obtained and were termed as heat stress responsive long non coding RNA (HS-LncRNA). Differential gene expression was studied between each sample (I,I+L and L) compared to the control condition. A total of 363 novel lncRNAs were identified with the probability >0.95 and ORF length >100 (Supplementary data). A total of 363 lncRNA that do not have sequence similarity to the known class of non-coding RNA were classified as heat stress lncRNA or “HS-lncRNAs”.

### Characterization of banana heat stress LncRNA

The characteristics and regulatory patterns of LncRNA were investigated in this study. The 363 newly identified heat stress related HS-LncRNA include 288 lincRNAs (long intergenic non coding RNAs), 71 antisense LncRNA, 5 sense lncRNAs (Figure 2). LincRNAs comprises the major part of the non-coding RNAs (79% of the banana lncRNAs). To identify the chromosome location of lncRNAs circos tool was used (Krzywinski et al. 2009). LncRNAs were observed to be evenly distributed on each chromosome with no obvious preferences of the location (Figure 3). This circular ideogram layout facilitates in identifying the position and orientation on a given genome. Similar patterns were observed in a study conducted on salt stress condition in *Medicago truncatula* where 140 lncRNAs were detected using NONCODE software (Zhao et al. 2016) by aligning it to *Arabidopsis thaliana* (Zhang et al. 2014).

**Figure 2:**
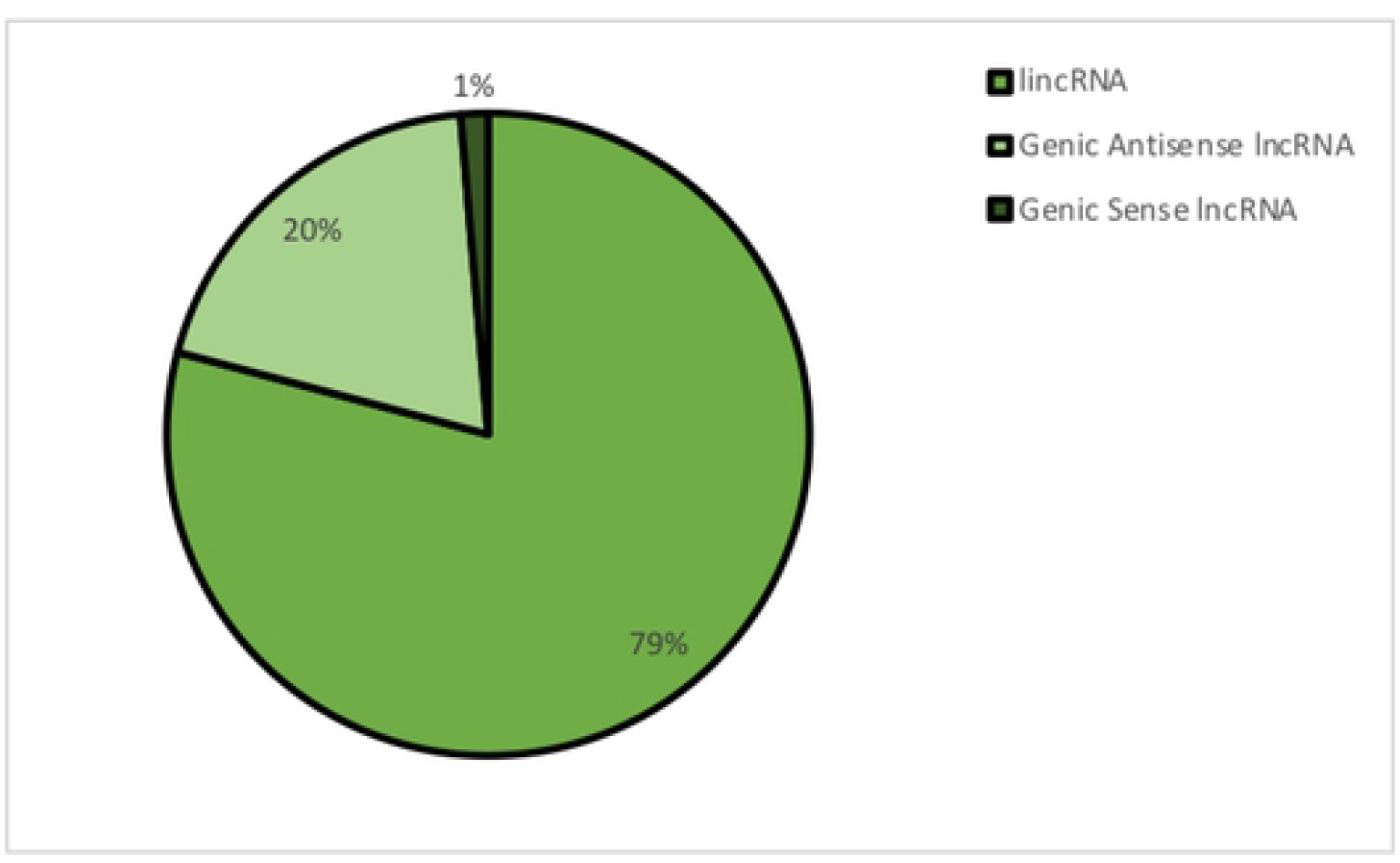
Classification of lncRNAs.

**Figure 3:**
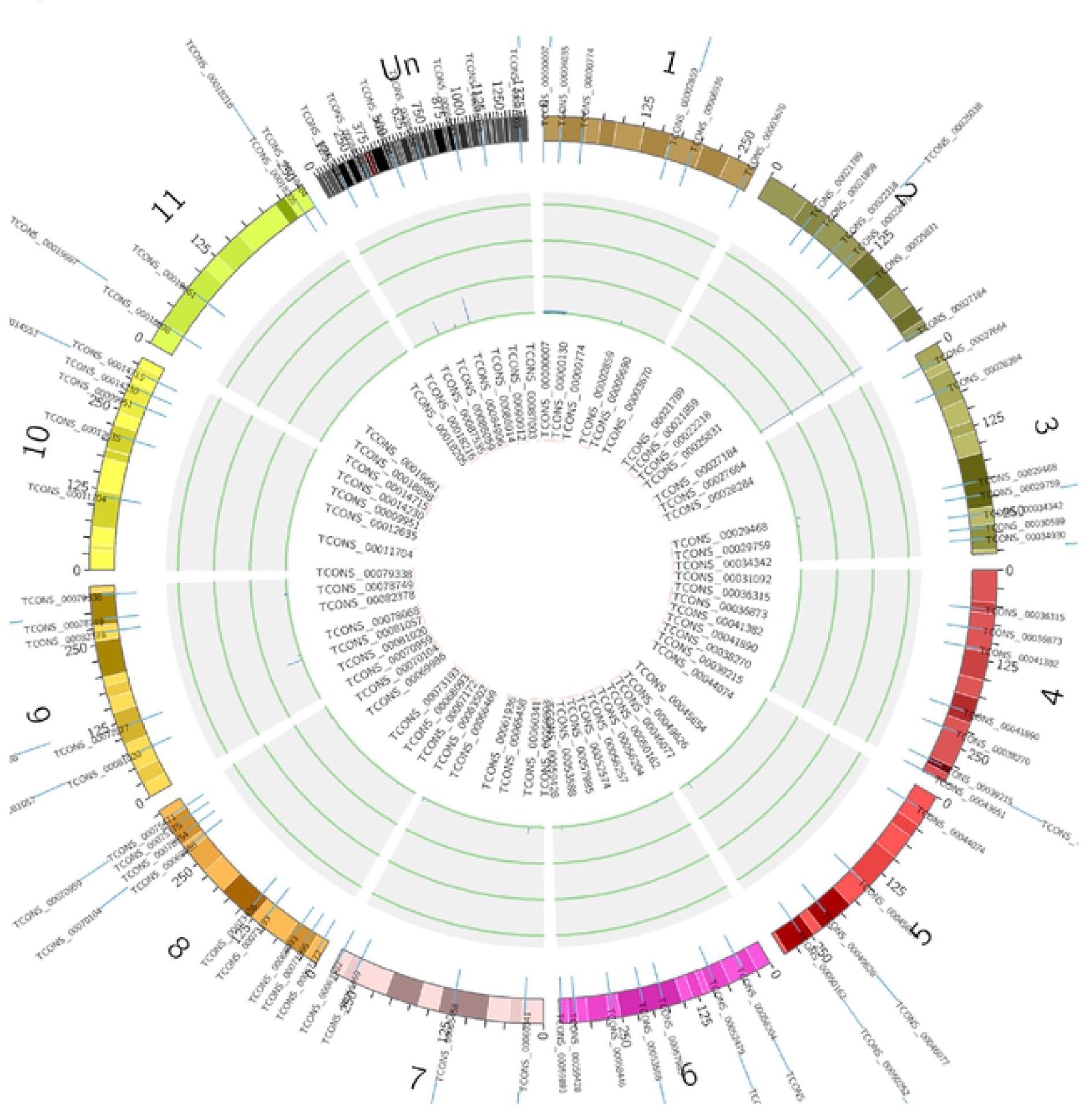
The expression levels of lncRNAs along the eleven chromosomes of banana. The plot consists of four concentric rings where each corresponds to a given sample.

### Functional analysis of heat responsive lncRNA

LncRNAs are known to regulate the neighbouring genes. These are also known to have *cis* roles. Therefore, to understand the potential roles of LncRNAs, Gene ontology analysis was carried out to identify the target genes using annocript software. Similarity search was done against BLASTX against TrEMBL/ UniRef and SwissProt, RPSBLAST against CDD profiles, BLASTN against Rfam and rRNAs. We identified significant enrichments (P<0.05) for all three components, for example biological process (GO:0006351, transcription, DNA-dependent; GO:0006412, translation) and cellular components (GO:0016021, integral to membrane; GO:0005634, nucleus; GO:0005886, plasma membrane) and molecular function (GO:0005524, ATP binding; GO:0003677, DNA binding; GO:0003677, structural constituent of ribosome; GO:0004672, protein kinase activity) (Figure 4). These findings suggest that genes are regulated by lncRNAs which might participate in different roles of transcription and translation, signal transduction, and metabolism in response to heat stress.

**Figure 4:**
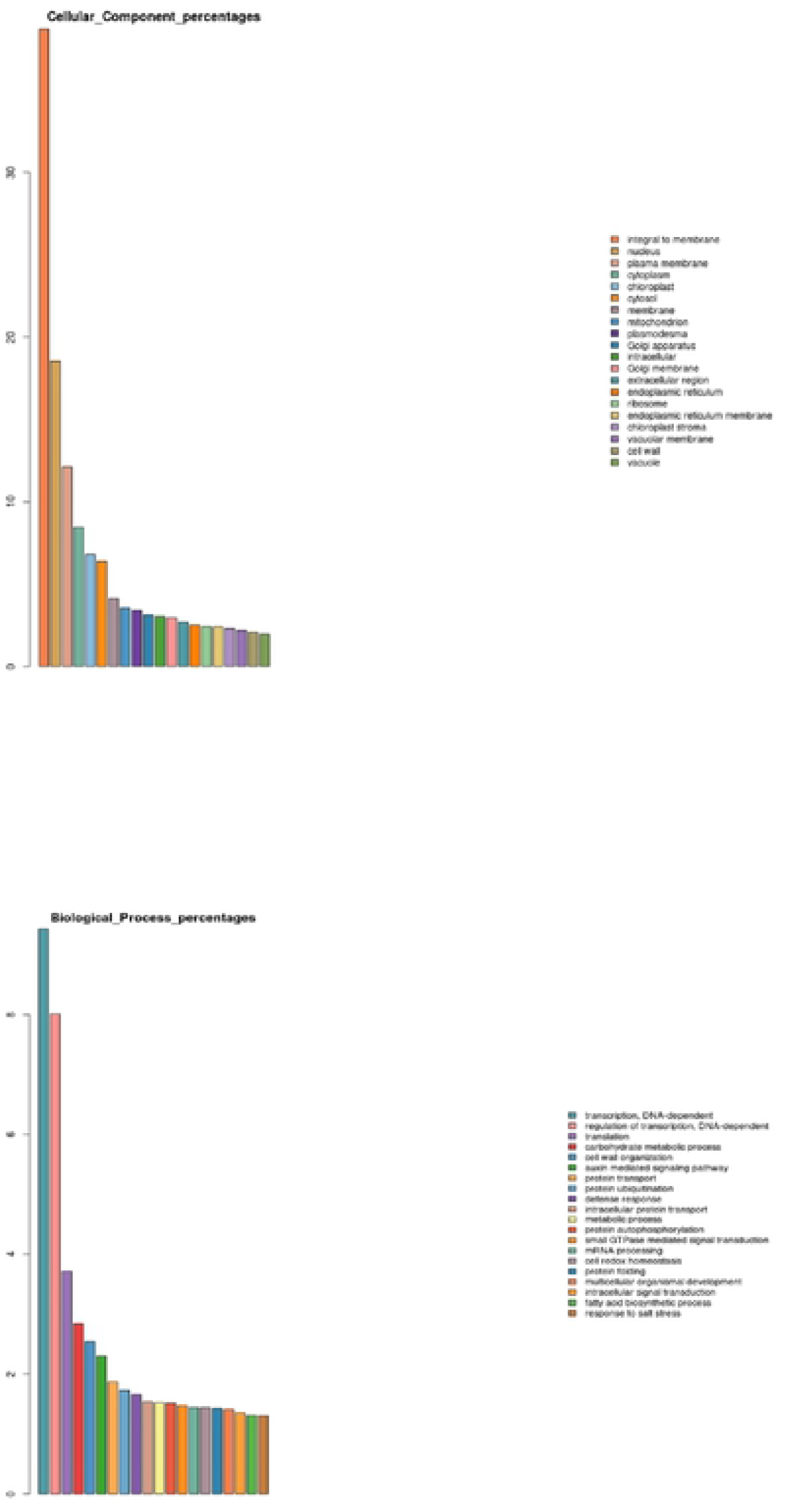
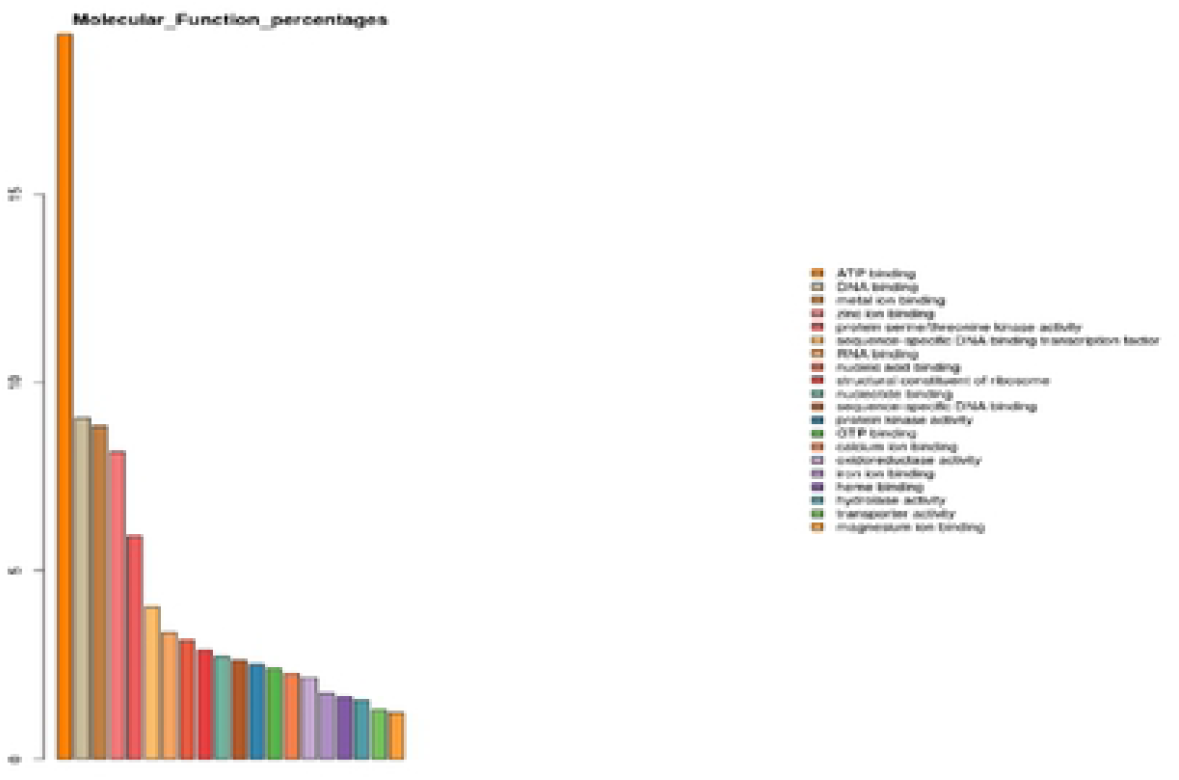
Gene Ontology of heat stressed leaves a) Cellular components b) Biological processes c) Molecular function.

LncRNA are known to regulate several biological processes and have diverse roles in development and also can regulate one to many protein coding genes (Liu et al. 2015). To understand this relationship of protein coding gene and LncRNA, Cytoscape was used to build an interactive plot or model network featuring the interaction of one to many genes cross talk (Figure 5). The interactions observed in the study had less than or equal to four nodes as shown in the figure 5. For example, the protein coding gene involved in the membrane integrity and other signal transduction were found to be regulated. Protein coding gene—N methyl transferase which was found in the interaction was regulated under heat stress, this suggests that these lncRNAs were involved in regulating the target mRNA/miRNA, i. e as lncRNA acts as a precursor molecule that regulates the other ncRNAs or mRNAs. Similar study was conducted in maize where the lncRNA and their interaction was studied (Zhao et al. 2014). One of the active models used in the study was mapping the cellular pathways responding to genetic perturbations and environmental stimuli (Schwikowski et al. 2003). The putative network built based on the protein coding and the vicinity of the lncRNA may not be very robust as the number of transcripts obtained were very few. Therefore, using this platform a span of several domains such as protein-protein interactions, transcriptional and post transcriptional changes can be visualized. Although, different data sets can be represented in different patterns and schemes but in the course of time they all reflect the same biological system and response.

**Figure 5:**
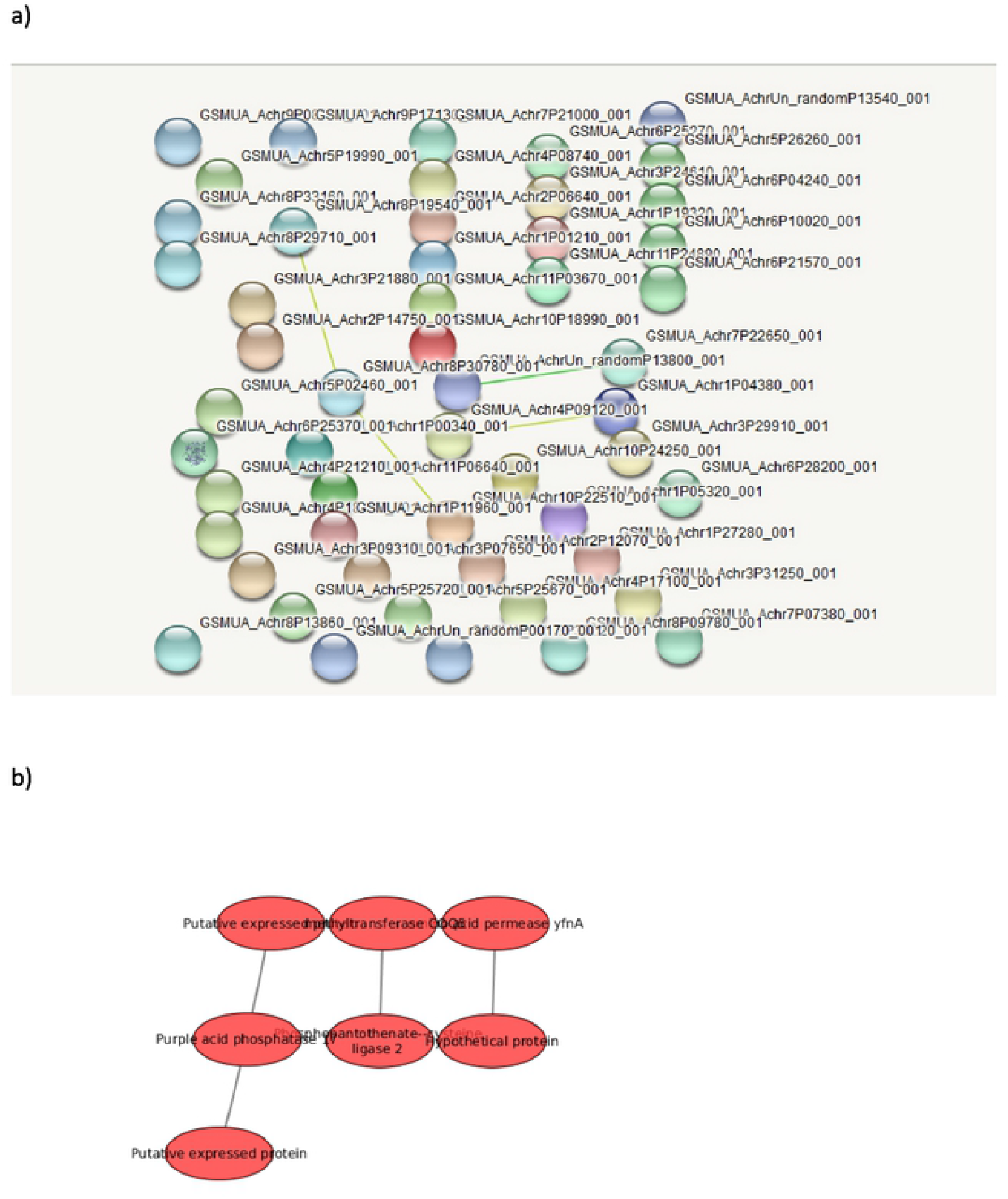
a) Interaction of lncRNAs b) Predicted protein coding genes using Cytoscape.

### LncRNAs as potential precursors of miRNA

miRNAs or microRNA are important class of non-coding RNA which are 20-22nt long in length. They are known for transcriptional and post transcriptional regulation of genes in plants (Chen et al. 2012). Recent studies have shown that major portion of lnRNA acts as a precursor of miRNA in many eukaryotes (Liu et al. 2012). In the present study, lncRNAs were aligned and miRNAs which were predicted using the FASTA sequences in psRNATarget software. A total of four miRNAs were found TCONS_00078956 precursor for miR8007b, TCONS_00027184 precursor for miR2083, TCONS_00075175 precursor for miR847 and TCONS_00022479 precursor for miR414. In recent reports, miR414 is known to be targeting AP2/B3-like transcriptional factor family protein, abscisic acid mediated signaling pathway (Xie et al. 2015). They are predicted to be participating in stress related signalling process. They also undergo several evolutionary changes due to large environmental or climate changes taking place due to which they become species specific or specific to abiotic stress.

### Differential expression of lncRNAs in response to heat stress

LncRNAs are actively involved in regulation of both biotic and abiotic stress (Shuai et al. 2014). In the present study, the normalized values for LncRNAs of control and heat stressed samples were calculated using FPKM (fragment per kilobase of transcript per million mapped reads). Of these DEGs, 21 that were differentially expressed in induction stress, 30 in I+L and 27 in lethal stress conditions were identified. Venn diagram depicts the abundance of transcripts at all three time points when compared to control (Figure 6). The study conducted on *Populus* reported a total of 504 drought responsive lincRNA (Shuai et al. 2014). 76 lncRNAs were identified in *A. thaliana* for abiotic stresses from full length cDNA library (Xie et al. 2015). In the present study, we identified four differentially expressed transcripts which were highly up regulated during induction stress, three during I+L and one during lethal stress condition. This suggests that banana HS-lncRNA mediated post transcriptional changes might occur at induction stress condition, which then gradually decreased during lethal stress. Heat map depicts the expression patterns under different stress conditions (Figure 7). It has been shown that TIR induction helps plants to gain acquired thermotolerance. Thus, acquired thermotolerance play a key role in regulating protein coding genes. To the best of our knowledge this is a first study on heat responsiveness of lncRNA in banana.

**Figure 6:**
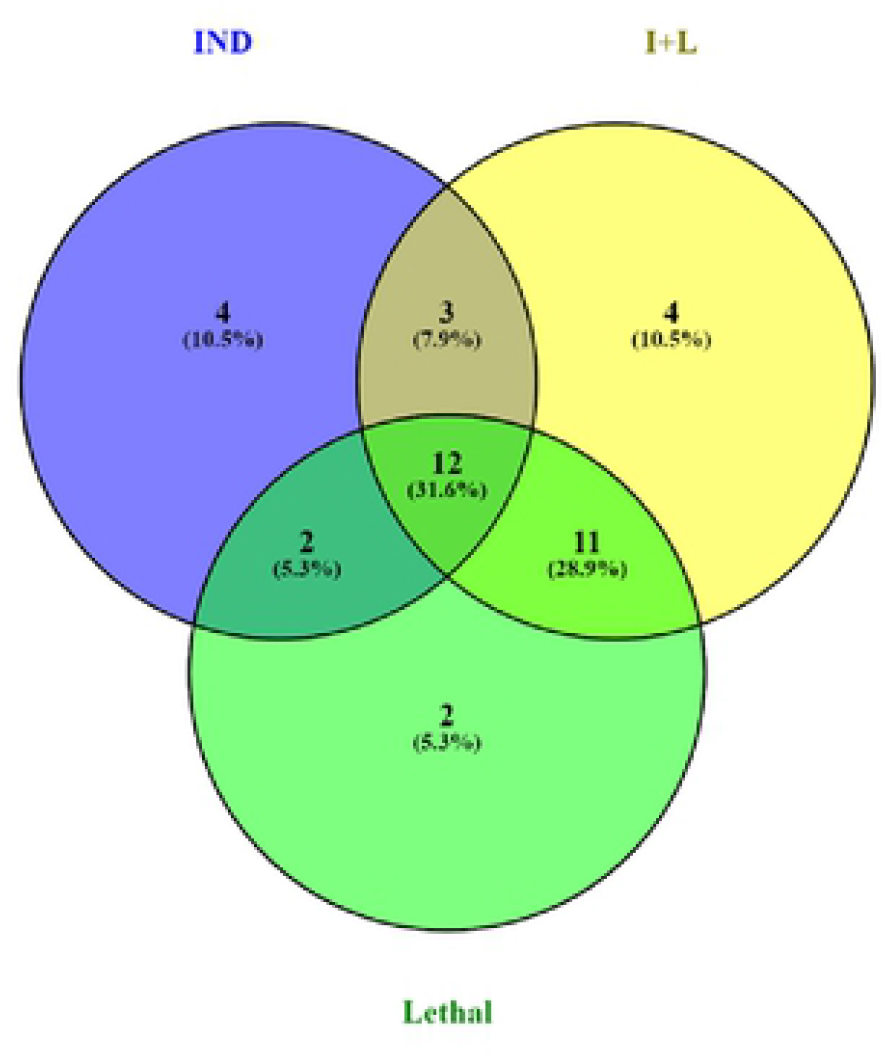
Venn diagram of DGE under heat stress.

**Figure 7:**
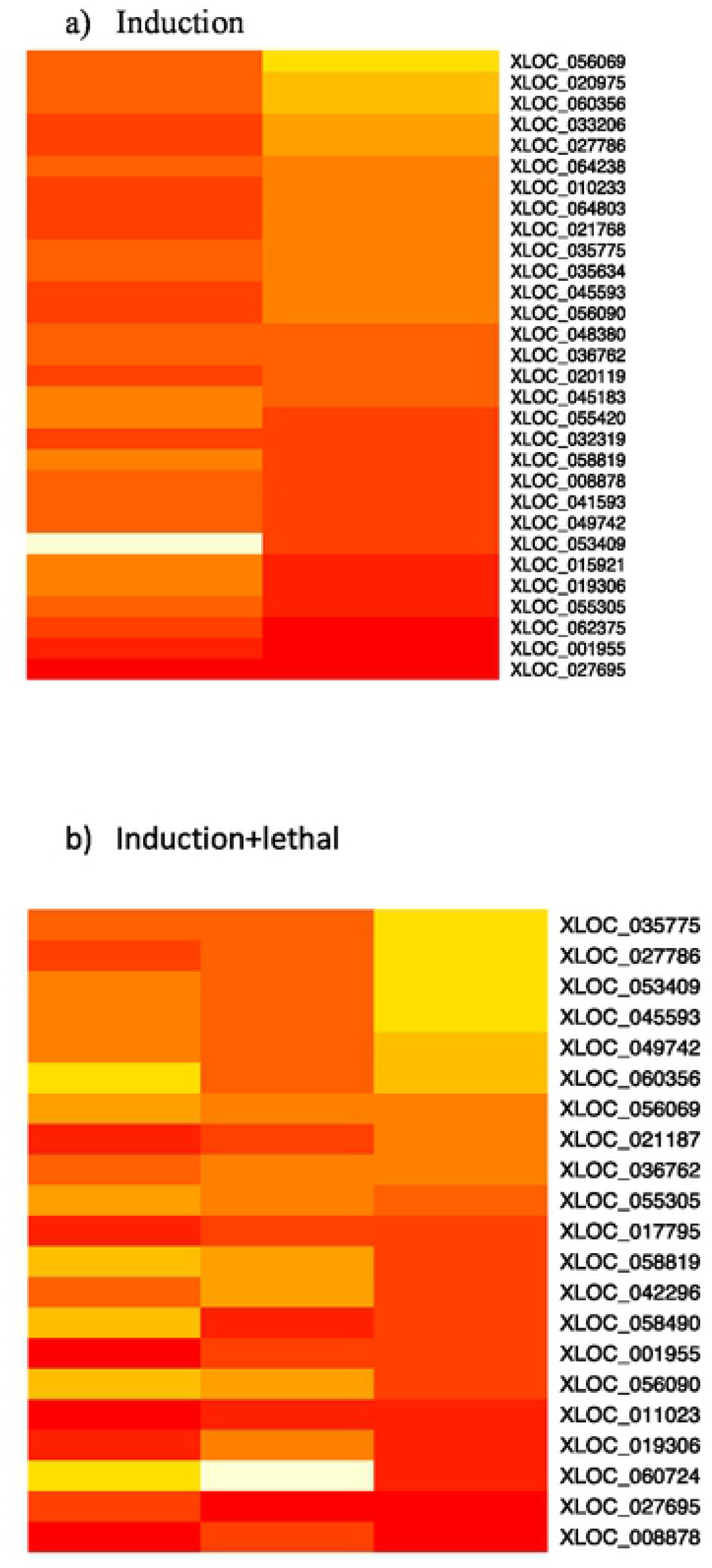
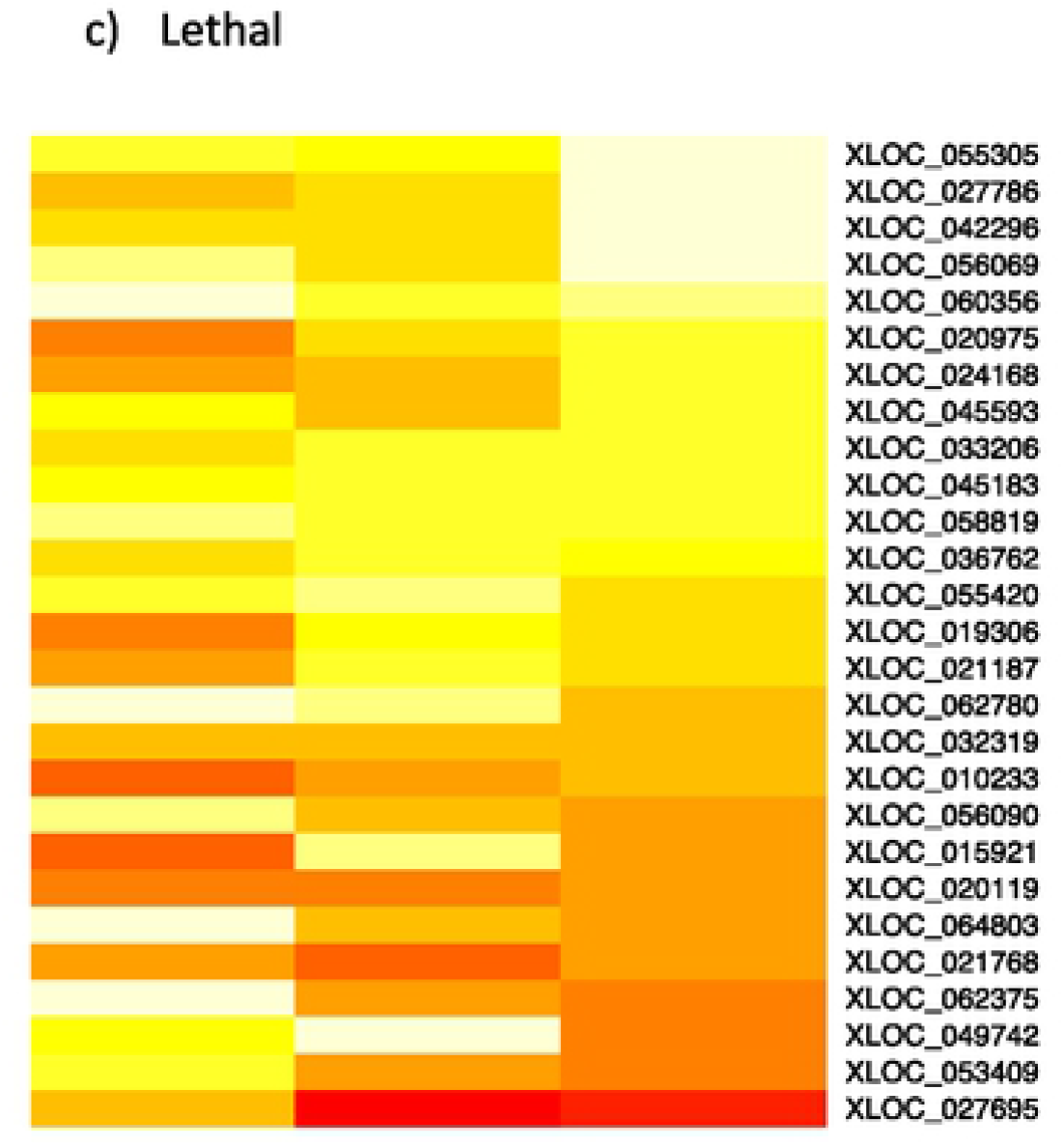
Heat Maps of a) Induction b) lnduction+lethal c) Lethal stress.

To validate the results obtained from RNA-seq, qPCR was used for investigating the expression of HS-lncRNA (TCONS_00075175, TCONS_00027184, TCONS_00078956, TCONS_00077877, TCONS_00051686) (Figure 8). As shown in the figure the expression patterns were compared between RNA-seq and qPCR which was found to be consistent with uniform patterns. The reliability with the qPCR data showed more expression when compared to RNA-seq with the correlation value of r^2^= 0.85 (P< 0.001).

**Figure 8:**
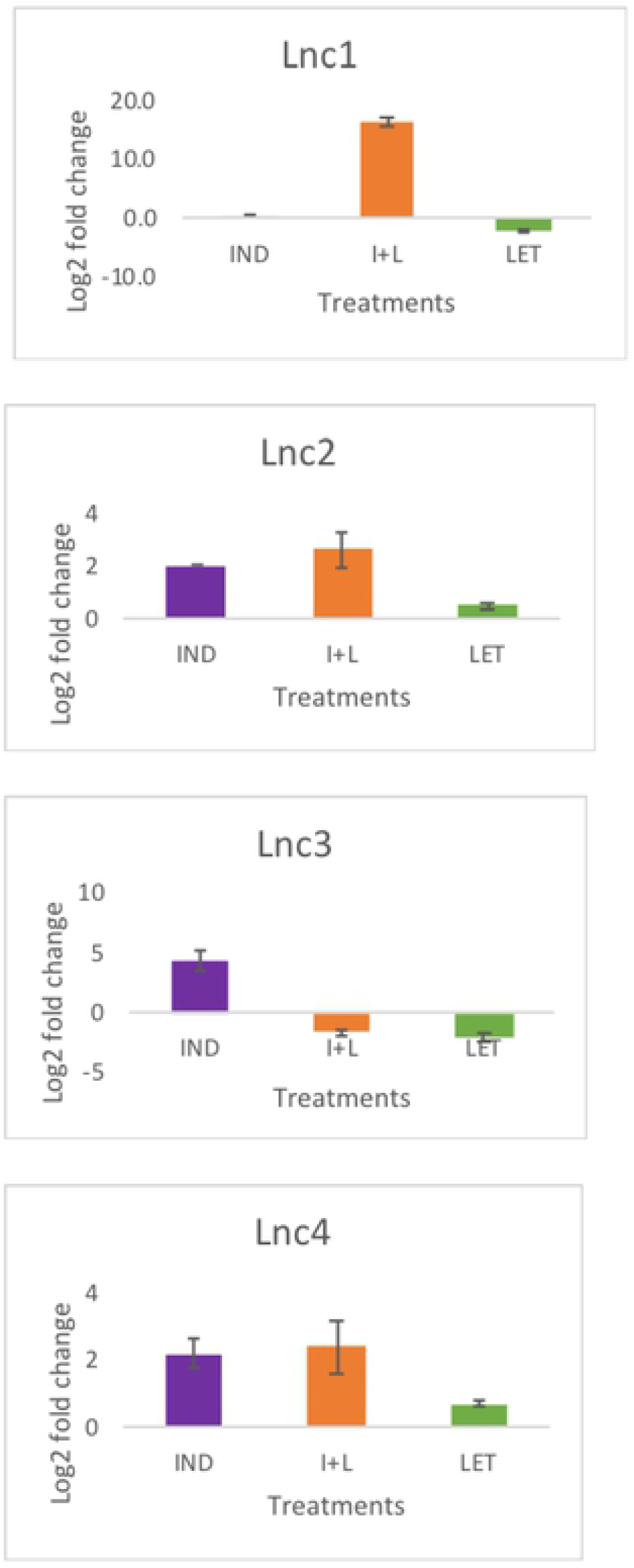
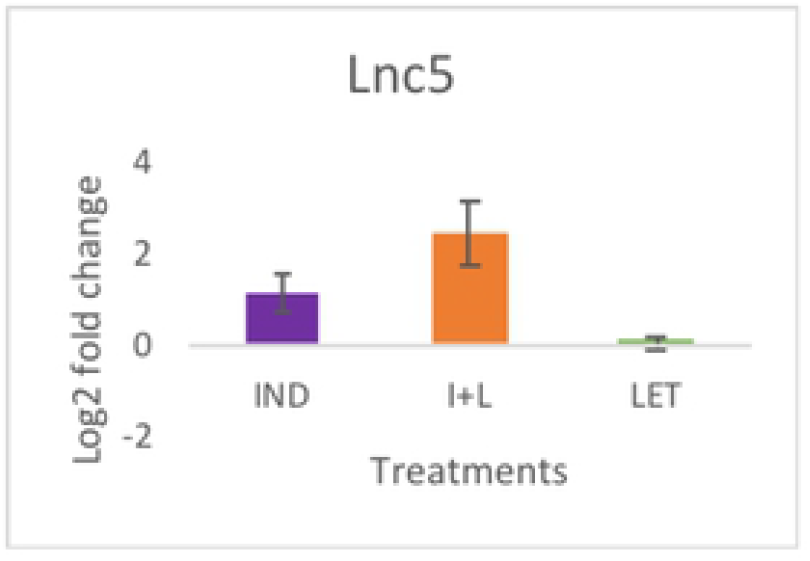
Expression levels of selected genes after Induction, Induction followed by lethal (I+L) and Lethal stress were examined by qPCR. The mRNA fold difference with respect to that of untreated (control). Bars show means ± SE (n = 3).

## Conclusion

In this study, we have identified 363 novel banana HS-lncRNA expressed during heat stress in banana. The LncRNAs were mapped to the banana chromosome and were found to be distributed evenly. Over to this a complex network pattern was studied between the protein coding genes and LncRNA. Further miRNA like miR8007b, miR414, miR2083, and miR847 were predicted where lncRNAs acts as precursor for target gene regulation including transcriptional and post transcriptional changes. All these evidences collectively suggest that LncRNA are presumably important in terms of stress conditions. However, little is known about the biological functions of lncRNAs in abiotic stress responses in plants. Moreover, lncRNAs are putative potent tools for plant improvement to enhance their resistance to abiotic stresses. Therefore, identification of heat stress-responsive lncRNAs, characterization of their functions and dissection of their regulatory networks can enhance our mechanistic understanding of plant response and adaptation to heat stress.

## Acknowledgments

The authors acknowledge the financial support from Indian Council of Agricultural Research (ICAR), New Delhi, India through National Innovations in Climate Resilient Agriculture (NICRA) project.

---

**Table S1:**
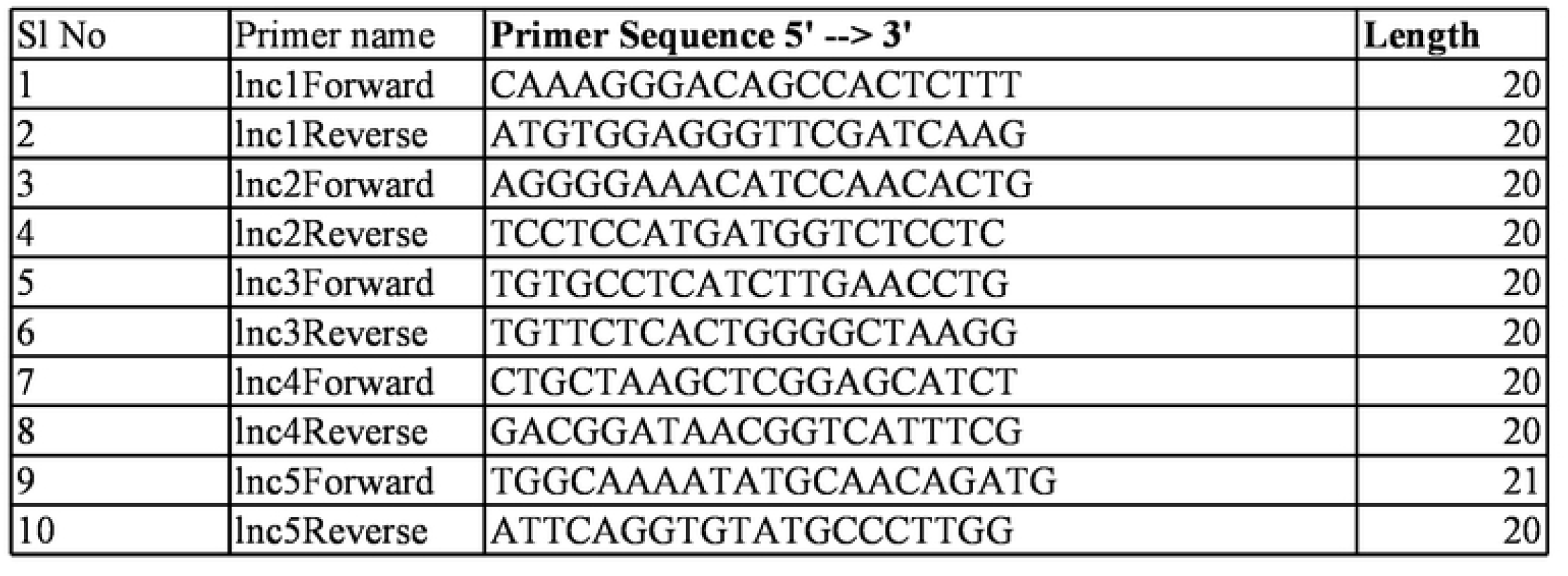
Primers used for qPCR

